# Conceptus attachment coincides with initiation of an anti-inflammatory cytokine profile in the pig endometrium

**DOI:** 10.64898/2026.05.01.722151

**Authors:** Günter P. Wagner, Thainá Minela, Alexandria Ross, Jan Engelhardt, Fuller W. Bazer, Gregory A. Johnson

**Affiliations:** Department of Ecology and Evolutionary Biology, Yale University, New Haven, CT 06520; Department of Evolutionary Biology, University of Vienna, A-1030 Vienna, Austria; Department of Veterinary Integrative Biosciences, Texas A&M University, College Station, TX 77843; Department of Animal Science, Texas A&M University, College Station, TX 77843

## Abstract

In eutherian mammals, blastocyst implantation is often associated with a quasi-inflammatory reaction in the endometrium, which is resolved with the establishment of the definitive placenta. This is understandable in the case of invasive placentation, since implantation entails a nidatory injury to the maternal tissue due to the invading blastocyst. Quasi-inflammatory processes have also been documented in pregnant pigs, even though the blastocyst only attaches to, rather than invades into, the endometrium of the uterus. In this study, we asked what processes in early porcine pregnancy lead to the resolution of attachment-associated inflammation. In generic wound healing the transition from a pro- to an anti-inflammatory state is caused by a corresponding transition from M1 to M2 polarized macrophages following efferocytosis by macrophages of apoptotic neutrophils. In order to determine whether this scenario applies to the pregnancy-related resolution of inflammation in the porcine uterus, we produced a series of bulk transcriptome samples spanning days (D) 13 to 25 of gestation. This time span corresponds to the transition from pre- to post-attachment stages of pregnancy. We found slower changes in the transcriptome between D20 and D25 than prior to D20, suggesting a turning point in pregnancy-related reprogramming. The turning point at D20 corresponds to the time of firm attachment of trophectoderm to uterine luminal epithelium and the cessation of IFNG signaling from the blastocyst. This transition coincides with increased expression of RNAs of genes implicated in resolution of inflammation and M2 polarization such as ARG1, MRC1/CD206, CD86, TGFb1 and IL10, as well as a significant increase in expression of HGPD, the enzyme that metabolizes prostaglandins. While immunoreactivity for ARG1 was found in putative macrophages in the sub-epithelial stratum compactum, other markers of M2 polarized macrophages were localized to non-immune cells: MRC1 was found on fibroblast-like stromal cells, CD86 on trophoblast cells, and IL10 in luminal and glandular epithelia. These results suggest that intrauterine immune regulation is decoupled from that of the rest of the body by engaging non-immune cell types as anti-inflammatory mediators during the peri-attachment period of pregnancy.

## Introduction

In pigs, which have epitheliochorial placentation, the morula enters the uterus, develops into a blastocyst, “hatches” from the zona pellucida to become a spherical blastocyst, and then elongates to form a conceptus (embryo and associated extra-embryonic membranes). Each conceptus within the litter eventually achieves a large surface area for contact between the trophoblast and endometrial luminal epithelium (LE) to facilitate uptake of nutrients from endometrial LE and glandular epithelia (GE) (Bazer and Johnson 2014, Johnson, Bazer et al. 2021). These elongating conceptuses secrete a series of steroids and proteins that are critical for the maintenance of pregnancy through the peri-attachment period (Bazer and Johnson 2014). Conceptus-derived estrogen is involved in blocking luteolysis of *corpora lutea* (CL) resulting in maintenance of high levels of secretion of progesterone necessary to support pregnancy, an event termed pregnancy recognition. Pig conceptuses’ elongation requires conceptus-derived IL1B. Conceptus-derived PGE2 and IFNG are hypothesized to be necessary for endometrial remodeling to support successful attachment of blastocysts and early placental development (Geisert, Bazer et al. 2024). In pigs, IFNG from conceptus trophectoderm is delivered to the endometrium from D13 to D20, at least in part, through extracellular vesicles (Cain, Seo et al. 2024). IFNG is required for resolution of inflammation in pigs (Geisert, Bazer et al. 2024). IFNG alters populations of T cells within the endometrium and upregulates or induces expression of multiple IFN-stimulated genes (ISGs) in the endometrial stroma (McLendon, Seo et al. 2020, Johns, Lucas et al. 2021, Johnson, Bazer et al. 2024, Sponchiado, Gu et al. 2025).

In eutherian mammals, the ancestral placenta was likely hemochorial and characterized by extensive invasion of the conceptus into the endometrium and decidualization of the endometrial stroma (Mess and Carter 2006, Wildman, Chen et al. 2006, Elliot and Crespi 2009). This condition is retained in humans in which uterine lacunae form around placental villi that fill with maternal blood providing for the exchange of nutrients, oxygen, hormonal signals, and waste products between the mother and the fetus (Cross, Werb et al. 1994, Norwitz, Schust et al. 2001). The intimate invasion of placental cells into the endometrial tissues necessitates that the endometrium modify the inflammatory response through the evolutionary origins of the decidual cells and specialized uterine immune cells (Mor, Cardenas et al. 2011, Erlebacher 2013, Chavan, Griffith et al. 2017, Stadtmauer and Wagner 2020). Pigs, on the other hand, have true epitheliochorial placentation as the endometrial LE remains intact throughout pregnancy and placental trophoblast cells attach to the endometrial LE but do not invade (Bjorkman 1973). This condition evolved secondarily from an invasive form of placentation, at the latest in the stem lineage of artiodactyls if not earlier (Elliot and Crespi 2009). As such, it has been predicted that pig conceptuses do not cause a nidatory injury and associated inflammation within the uterus (Joyce, Burghardt et al. 2008). Nevertheless, it has been shown that the peri-attachment period of pigs is associated with a substantial quasi-inflammatory and adaptive immune reaction (Samborski, Graf et al. 2013, McLendon, Seo et al. 2020, Johns, Lucas et al. 2021). This quasi-inflammatory process is thus likely an evolutionary carryover from hemochorial ancestors retaining those parts of the inflammation process necessary for endometrial remodeling in preparation for continued gestation.

For decades, the immunology of pregnancy was thought to follow principles of transplantation biology, with the maternal immune system responding to the fetus as it would to a foreign graft. This view arose from the recognition that the conceptus must avoid immune recognition by the maternal adaptive immune system to prevent rejection (Medawar 1953, Medawar 1961). However, this focus has been unfortunate in that most investigators thought that the events of conceptus attachment and implantation require an anti-inflammatory immune environment within the endometrium (Wegmann, Lin et al. 1993). As a result, it has been overlooked that the process of implantation contends with inflammation caused by nidatory injury, also known as the inflammation paradoxon (Chavan, Griffith et al. 2017), that must be resolved before the inflammation can damage the conceptus. In this study, we hypothesized that there is a signature of processes leading to resolution of inflammation during the peri-attachment period of pregnancy in the pig. We find that this is indeed the case, but, surprisingly, with different cellular players than those responsible for the resolution of inflammation during wound healing.

## Results

During the peri-attachment phase of porcine pregnancy, there is a documented active inflammatory and immune response (Samborski, Graf et al. 2013, McLendon, Seo et al. 2020). However, later stages of pregnancy are characterized by an immunologically quiescent endometrial environment. In order to gain insights into the mechanisms behind this transition we conducted a time series of bulk RNAseq experiments and followed up with immuno-localization of selected proteins based on the transcriptomic results.

### Dynamic endometrial transcriptional profile during the peri-attachment phase of pregnancy

In order to investigate gene expression and cell composition changes in the porcine endometrium during the peri-attachment period of pregnancy we sequenced the bulk transcriptomes from 15 samples collected between day 13 (D13) and D25 of pregnancy that includes both pre- and post-attachment stages of pregnancy (Johnson, Bazer et al. 2021). These include D13 (N=3), D15 (N=4), D17 (N=2), D20 (N=3) and D25 (N=3). RNA was extracted, converted into cDNA that was sequenced, and the reads mapped to the porcine genome (Sscrofa 11.1). RNA abundance was quantified as transcripts per million (TPM) (Wagner, Kin et al. 2012). Data were pre-processed by eliminating non-expressed and very lowly expressed genes (max_across samples_(TPM)<3) to avoid data distortion from a large number of non- and very lowly-expressed genes. The reduced dataset included 15,440 genes.

To test for internal consistency of expression profiles, Spearman correlation coefficients between all pairs of samples were calculated and represented in a heatmap (Figure 1A). All samples from the same day of pregnancy are internally consistent except for two samples from D17. One of the D17 samples (animal ID = G8) appears to be more similar to D15 samples and the other (animal ID = G21) more similar to D20 samples. This variation likely suggests that there is a fast transition between D15 and D20 making samples from D17 more variable. The remaining samples represent two blocks of internally correlated samples, D13 and D15, as well as D20 and D25. This pattern suggests a faster change in gene expression between D15 and D20 than between D20 and D25. This is consistent with the fact that these three sample days, D15, D20 and D25, are separated by 5 days. However, the earliest sample, D13, is only two days earlier than the D15 samples, and thus the similarity between D13 and D15 samples might be due to the close time points in pregnancy rather than reflecting a slower rate of change between D13 and D15 compared to D15 to D20.

**Figure 1:**
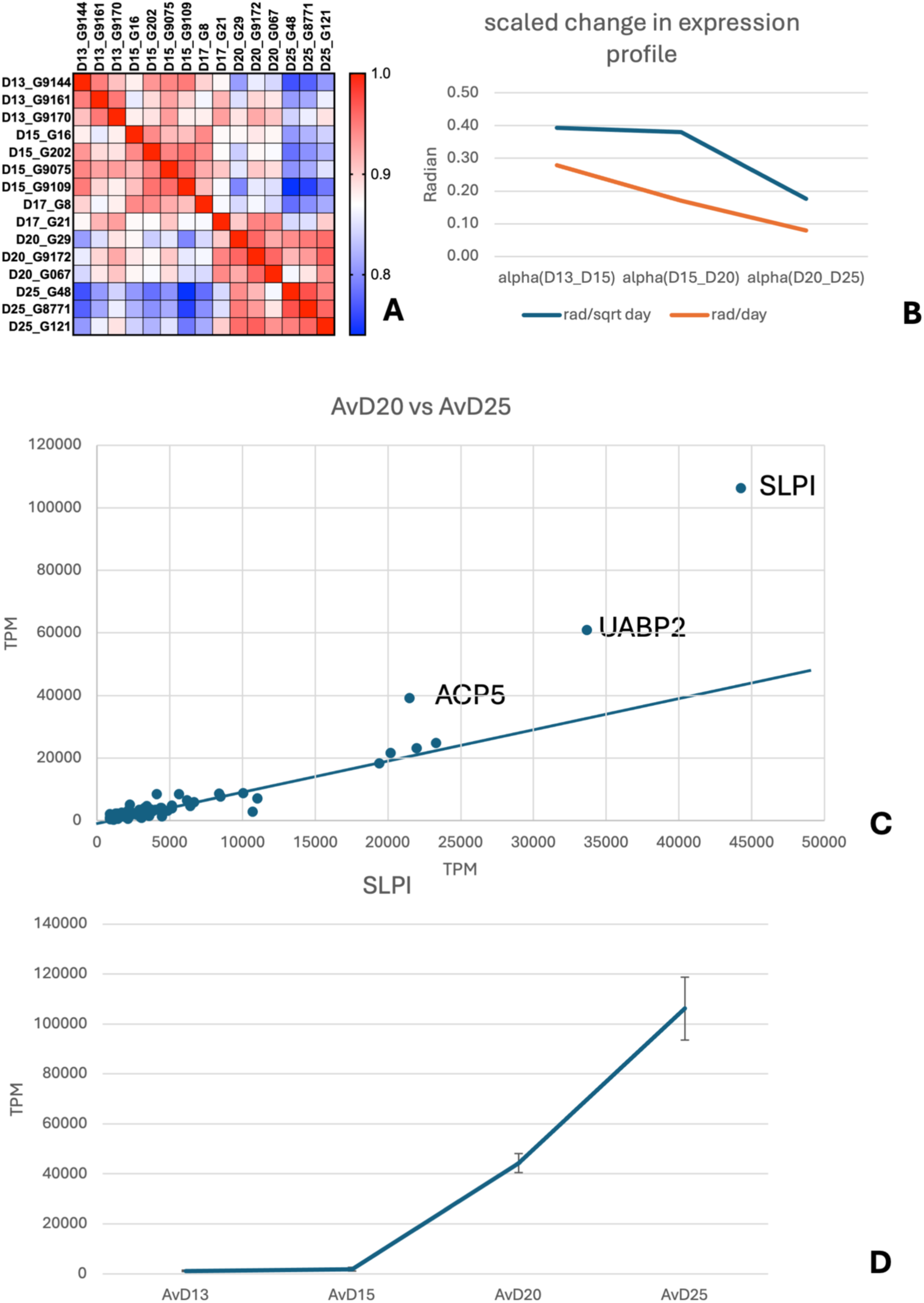
Transcriptome dynamics in the endometrium during the peri-attachment phase of pregnancy in pigs. **A**) Heatmap of transcriptomic similarity for 15 sequenced samples representing D13, D15, D17, D20, and D25 of pregnancy. Transcriptome similarity was quantified by Spearman correlation coefficients, that is, rank order correlation, to avoid the effect of a few highly expressed genes. Note that the transcriptomes from D13 and D15 are more similar to each other than those for D20 and D25, and these are more similar to each other than those from the earlier stages. **B**) Rate of change in the transcriptome was estimated by the angle between average gene expression profiles between D13 and D15, D15, and D20 as well as between D20 and D25. The average rate per day was estimated using two methods: dividing by the number of days, which assumes that changes happen in a straight line in gene expression space, and the second, which assumes that direction of change from day to day is random, i.e., the new direction is orthogonal to the direction in the previous stage. **C**) Comparison of average expression of the 50 most highly expressed genes on D20 and D25. Most of the genes have similarly expressed on D20 and D25 except for three genes: *SLPI, UABP2* and *ACP5*, which are much higher expressed on D25 than D20. **D**) expression dynamic of *SLPI*, secretory leukocyte protease inhibitor for which expressed is low up to D15 and then increases coincidentally with development of firm conceptus–LE attachment.

To estimate the rate of change per day we calculated the angle between the gene expression vectors of consecutive pregnancy days (ignoring D17 samples) and scaled the amount of change either per day or per the square root of the number of days between time points (see M&M for justification of square root scaling, Figure 1B). The results suggest that the overall rate of change per day in the gene expression profile is greater between D13 and D15 and D15 and D20 than the rate of change between D20 and D25, suggesting a consolidation of the gene expression profile following attachment. However, inspection of the length of the expression vectors revealed a substantial increase from D20 to D25. The length of the expression vector is related to the amount of variation among genes, i.e. a greater length suggests a steeper gene expression profile in samples with longer expression vectors, given that the sum of TPM values is constant per definition. To further investigate this effect, we plotted the average TPM values from D25 over those from D20 for the 50 highest expressed genes (Figure 1C). This plot revealed overall similar gene expression levels on D20 and D25, except for three genes that are substantially more highly expressed on D25 than on D20. These are secretory leukocyte protease inhibitor (*SLPI,* ENSSSCG00000039573) (Figure 1D), uteroferrin-associated basic protein 2 (*UABP2*, ENSSSCG00000022251) (Suppl. Fig. 1A) and tartrate-resistant acid phosphatase (*ACP5*, ENSSSCG00000032282) (Suppl. Fig. 1B). The proteins coded for by these genes are well-known components of the histotroph of the pig endometrium (Fazleabas, Bazer et al. 1982, Bazer and Roberts 1983, Fazleabas, Geisert et al. 1983, Roberts and Bazer 1988, Bailey, Dunlap et al. 2010, Steinhauser, Bazer et al. 2017). Specifically, SLPI expression begins on D20 and increases to D25, but is low up to D15 (Figure 1D). These results indicate that these genes undergo a steep increase in expression after conceptus attachment, unlike most other highly expressed genes. This expression dynamic is interesting because of the potential role of protease inhibitors expressed by the LE in preventing blastocyst invasion into the uterus (Fazleabas, Bazer et al. 1982). Outside the uterus, the pig blastocyst is known to be invasive (Samuel 1971, Samuel and Perry 1972), and the lack of invasiveness is thus due to endometrial resistance (Fazleabas, Bazer et al. 1982).

### Abundance of RNAs of immune regulatory genes between pre- and post-attachment stages of pregnancy

To compare abundances of RNAs from candidate genes before and after attachment we pooled samples from D13 and D15, including D17 gilt number 8 (G8) into a pre-attachment sample (N=8) and D20 and D25 including D17_G21 into a post-attachment sample (N=7). We first investigated differences in the expression of genes related to immune function, with changes considered significant at p < 0.05, suggestive if 0.05 < p < 0.1, and nominally different if the means are numerically different but p > 0.1.

The abundance of the pan-leukocyte marker *CD45* protein tyrosine phosphatase receptor type C (*PTPRC*) is greatest during the post-attachment state (3.18x increase, p=9.3x10^-3^), suggesting an increase in the abundance of leukocytes (Figure 2A); however, it is difficult to assign this increase in leukocyte marker gene expression to any particular type of leukocyte. There is no increase in the abundance of the macrophage marker *CD68*, a nominal decrease in the neutrophil gene elastase neutrophil expressed (*ELANE)*, and nominal increases in the Th2 marker *GATA3* and Treg marker fork head box protein P1 (*FOXP1)*, as well as the other T-cell markers including *CD3, CD4* and *CD8*, but none of them are statistically significant (NK cells are discussed below) (Figure 2A). It is thus likely that the significant increased abundance of the pan-leukocyte marker *CD45* mRNA is due to small contributions from multiple leukocyte cell types, or higher CD45 expression in constant leukocyte populations.

**Figure 2:**
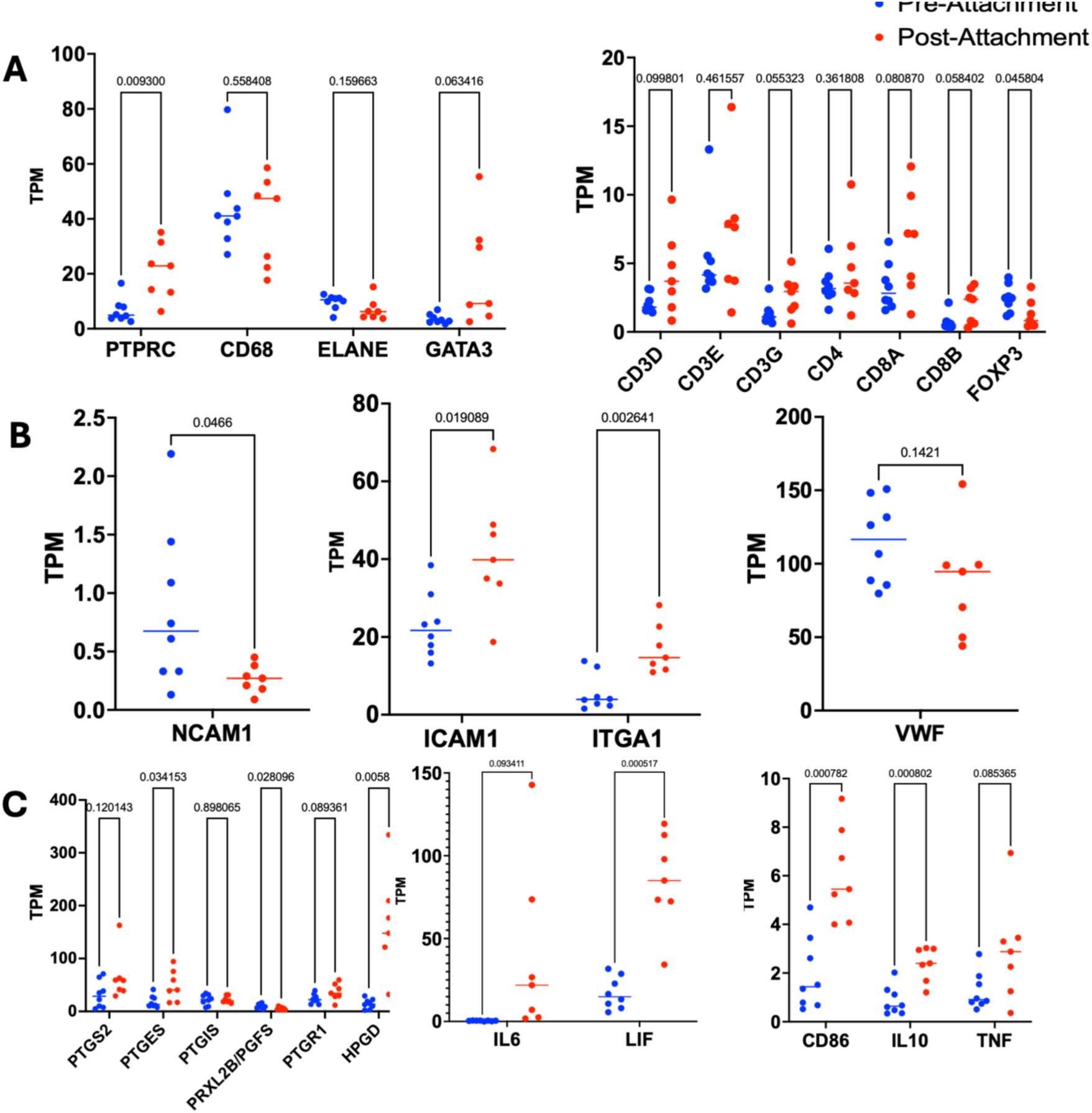
Comparisons of endometrial expression of selected immune-related genes in pre-attachment, and post-attachment stages of pig conceptuses. **A**) Expression comparisons of leukocyte marker genes. **B**) Changes in expression of marker genes for NK and endothelial cells. **C**) Changes in expression of genes related to prostaglandin metabolism (left panel), and inflammation-regulating cytokines (two right panels).

In the human endometrium, decidual natural killer cells (dNK) are characterized as CD56^bright^ and CD16^dim^. Expression of *CD56* mRNA (aka NCAM1) was low overall in pig samples (<2TPM) and even lower in post-attachment than pre-attachment samples (0.3x decrease, p=0.0466) (Figure 2B). In contrast, expression of two markers of tissue resident NK cells was substantial and greater in post-attachment than pre-attachment samples: intercellular adhesion molecule 1 (*ICAM1;* aka *CD54*; 2.2x increase, p=0.019) and integrin subunit adhesion molecule 1 (*ITGA1;* aka *CD49a*; 2.98x increase, p= 2.6 10^-3^). Since ICAM1 is also abundant in endothelial cells, this increase in post-attachment samples could be due to the expansion of blood capillaries. An increase in the size of capillaries in the vasculature immediately beneath the endometrial LE has been documented (Seo, Li et al. 2020) and H&E staining has demonstrated increasing accumulation of red blood cells within the blood vessels of the stratum compactum stroma of the endometrium at attachment sites in pigs on D12 and D24 of gestation (Johnson, Seo et al. 2023). To test this interpretation, we also determined the abundance of the endothelial marker *Von Willebrand Factor* (*VWF*) RNA and found no significant change in *VWF* RNA abundance, and even a nominal decrease (Figure 2B). Together, these results suggest that there is an increase in tissue resident NK cells during the attachment phase of pregnancy in pigs.

### Increased expression of prostaglandin catabolic gene 15-hydroxyprostaglandin-dehydrogenase (HPGD)

There are signs that the prostaglandin system is more active prior to than after conceptus attachment (Figure 2C). We found a nominal two-fold increase (p=0.12) in the abundance of prostaglandin endoperoxidase synthase 2 (*PTGS2*, aka *COX2)* RNA. In contrast, there was a significant increase in PGE2 synthase RNA (*PTGES*; 2.6x increase, p=0.034) and a significant decrease in PGF2 synthase mRNA *Peroxiredoxin like 2B* (*PRXLS2B*) (0.55 TPM ratio, p=0.028) from pre-attachment to post-attachment phases. Most notable, the primary prostaglandin catabolic enzyme, 15-Hydroxyprostaglandin dehydrogenase, was significantly increased post-attachment (*HPGD*; 11.4x increase, p=5.8 10^-3^). This increase is consistent with the work of Seo and collaborators, who reported an increased expression of *HPGD* mRNA from D30 to D90 and then a decrease in expression to D114 when parturition occurs (Seo, Choi et al. 2014). Overall, these results suggest that conceptus attachment is associated with downregulation of prostaglandin signaling, given that the strongest increase in RNA expression was observed for HPGD, the key catabolic enzyme for prostaglandin inactivation, despite maintained and even increased expression of prostaglandin synthases.

### M2-like gene expression signature post-attachment

During the peri-attachment phase of pregnancy, in pigs an increase in inflammatory processes and immune cell types has been documented (McLendon, Seo et al. 2020). Thus, we investigated whether the resolution of attachment-related inflammation is caused by activation of classical alternative macrophages (Serhan and Savill 2005, Sugimoto, Sousa et al. 2016, Panigrahy, Gilligan et al. 2021), i.e. higher expression of markers for the inflammatory M1 macrophages pre-attachment and higher expression of M2-linked markers post-attachment. Changes in abundance of M1 markers were not consistent with a model of macrophage activation during the peri-attachment stage of pregnancy (Suppl. Figure 2).

Three generic M2 macrophage markers, Mannose Receptor C-Type1, MRC1 (aka CD206), Arginase 1 (ARG1), and Interleukin 10 (IL-10), showed significantly higher RNA expression post-attachment (Figure 2C and 3A). MRC1 has been reported to be expressed on macrophages and immature dendritic cells as well as some fibroblasts and keratinocytes (Gordon 2003). MRC1 has anti-inflammatory potential in that it facilitates endocytosis and neutralization of mannose-decorated cytokines and hormones and thus has anti-inflammatory potential. In the present study, MRC1 mRNA expression was lowest on D15 and increased through D20 and D25. Immunoreactivity (ImR) for MRC1 was detected in fibroblast-like cells of the endometrial stroma (Figure 3B and C) and placental compartment (Figure 3D). The MRC1-positive cells in the placenta were larger and rounder than the cells in the maternal stroma, suggestive of macrophage identity, but to our knowledge, the identity of these cells is unknown, but likely are homologous to Hofbauer cells from the human placenta. Some MRC1-positive macrophage-like cells were also detected in blood vessels (Figure 3E). ARG1 mRNA expression was not detectable on D13 and D15 (max(TPM)=0.18, av(TPM)=0.11), but was detectable at low and highly variable levels on D20 and D25 (min(TPM)=0.3, av(TPM)=1.6). Consistent with the variable expression of ARG1 mRNA, detection of ARG1 ImR was variable, but ARG1 ImR was detected on macrophage-shaped cells in the sub-epithelial stratum compactum on D20 (Figure 3F). The localization of IL-10 ImR will be discussed later.

**Figure 3:**
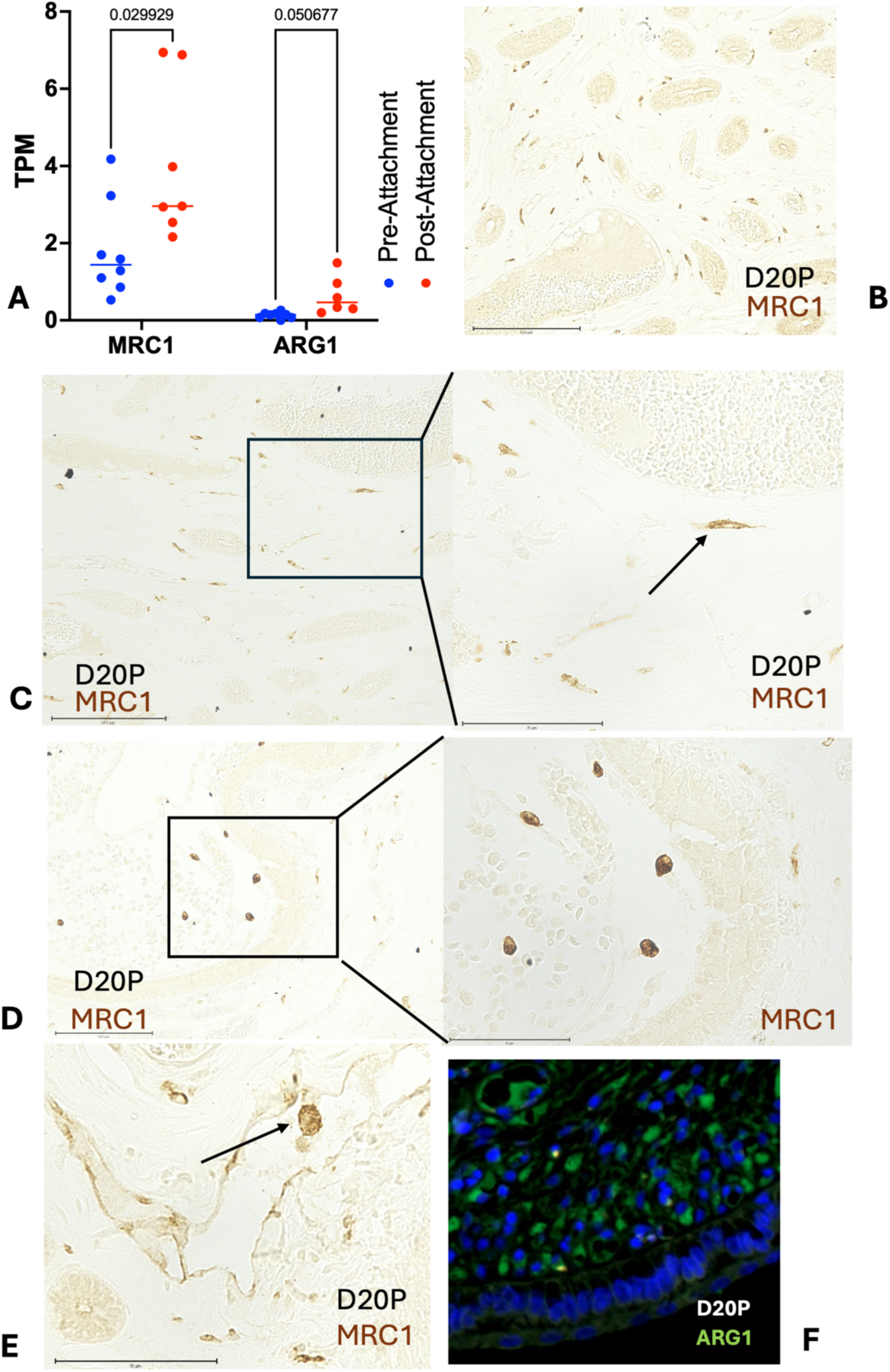
Expression of mannose receptor MRC1 (aka CD206) and arginase 1 (ARG1), both M2 macrophage markers: **A**) RNA expression levels in pre- and post-attachment stages of porcine pregnancy. **B**) Low magnification pattern of MRC1 immunoreactivity at D20 near endometrial glands showing cells with fibroblast-like morphology. **C**) Higher magnification of MRC1 reactive cells on D20 of porcine pregnancy. **D**) MRC1-positive cells in the placental compartment of the fetal-maternal interface on D20. **E**) MRC1-positive putative macrophages within a blood vessel in the endometrium. **F**) ARG1 immunoreactivity on putative macrophages in the stratum compactum stroma of the endometrium.

It is noteworthy that there were substantial increases in abundance of mRNAs for the M2b/Mreg marker genes, *CD86, IL10,* and *TNF*, post conceptus attachment (Figure 2C and 3A). The abundances of both *CD86* and *IL10* RNAs strongly increased post-attachment (*CD86*, 3.1x increase, p= 7.82 10^-4^; IL10, 2.8x increase, p=8.02 10^-4^). There was also a suggestive trend toward increased TNF and IL6 mRNA abundances (TNF, 3.2x increase, p=0.085; IL6, >100x increase, p=0.093), but with highly variable abundances among post-attachment samples, leading to low statistical support. The increase in Leukemia Inhibitory Factor, a member of the *IL6* cytokine family, *LIF*, (4.9x increase, p=5.17 10^-4^) post-attachment was remarkable and consistent with results from previous studies (Modric, Kowalski et al. 2000, Blitek, Kaczmarek et al. 2010).

### CD86 protein is expressed by trophoblast cells and appears to be transferred to the luminal epithelium

To assess whether the putative M2/Mreg signature found in the bulk transcriptome is in fact caused by an M2/reg polarization of uterine macrophages we performed immunolocalization for CD86 on uteroplacental sections from D15, D20 and D25. CD86 often serves as a cell surface ligand on antigen-presenting cells, such as macrophages and dendritic cells, as well as on some B-cells. CD86 engages with either CD28 or CD152 (aka CTLA-4) on T-cells. In D15 uteri, ImR for CD86 was found in the cytoplasm of trophoblast cells not yet attached to the LE, and in some specimens, it was localized to the apical segment of the trophoblast cells attached to the LE (not shown). On D20, intense CD86 ImR was localized at the apical segment of the trophoblast cells in areas of placental-uterine attachment (Figure 4A). On D25, CD86 ImR was found in the apical segment of the cytoplasm of uterine LE (Figure 4A, lower right image), a change reminiscent of trogocytosis that has been observed during interactions between macrophages and T-cells (Joly and Hudrisier 2003). In addition, CD86 ImR was also found in the perivascular cells of putative arterioles (Suppl. Figure 3), but no CD86 ImR was identified in any other cells within the stromal compartment of the endometrium during any stage of pregnancy examined.

**Figure 4:**
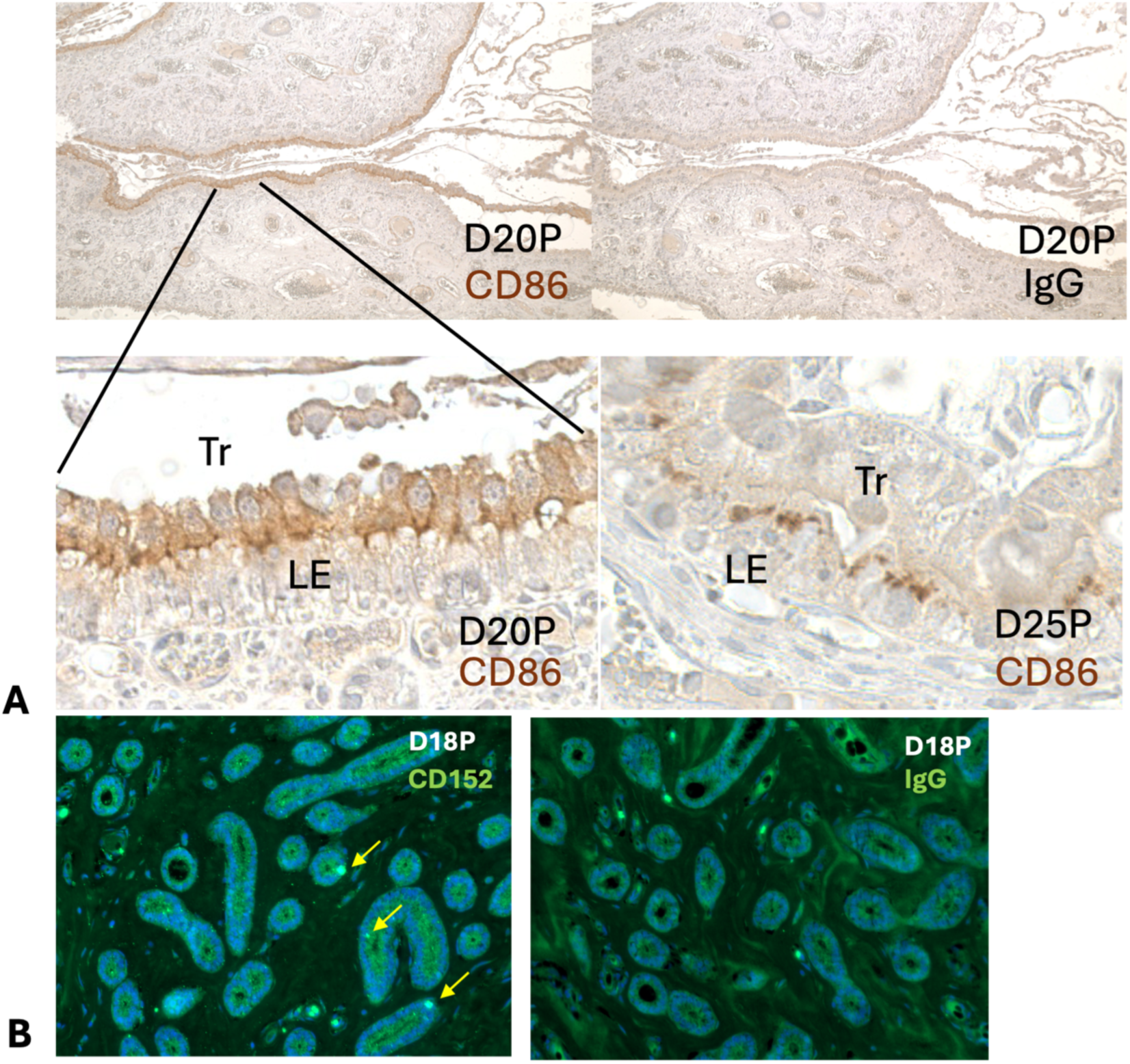
Expression of CD86 and CD152: **A**) CD86 expression on D20 and D25 of porcine pregnancy. Top left low magnification image of CD86 staining; top right negative control; bottom left magnification of part of the LE from the top left image showing CD86 immunoreactivity on the apical side of the trophoblast epithelium; bottom right showing CD86 immunoreactivity in the apical cytoplasm of the LE in apparent trogocytosis. **B**) CD152 expression on D18. Left image shows immunoreactivity within the glandular epithelium, possibly intra-epithelial T-cells; right image, IgG control.

The presence of CD86 at sites of attachment between trophoblast cells and endometrial LE on D20 and its putative trogocytosis into LE cells suggests that LE cells express a receptor for CD86 on their apical surface, which has yet to be identified. Two potential canonical CD86 receptors are known: CD28 and CD152/CTLA-4. CD28 activates T-cells and CD152 is implicated in inhibiting activated T-cells. Transcripts of both genes were detected in our RNAseq data, but at very low abundances. To test whether trophoblast CD86 engages with one of these receptors we immunostained for CD152 and CD28. We found strong CD152 ImR in single, round cells embedded within the endometrial GE, which could be intra-epithelial T-cells (Figure 4B). However, in the LE CD152 ImR was not consistently identified, in spite of multiple attempts, suggesting that CD152 is unlikely the receptor of the trophoblast CD86. Similarly, attempts to find ImR for CD28 on endometrial epithelia were unsuccessful. Hence, at this point, we cannot clarify the nature of the CD86-mediated interaction between the trophoblast and the LE, if any.

### IL10 protein is expressed in the epithelia of the endometrium

IL10 is a key cytokine produced by type 2 regulatory macrophages responsible, to a large degree, for their anti-inflammatory activity. IL10 RNA shows increased expression in post-attachment samples (2.6x increase, p=8 10^-4^; Figure 2C). We investigated the cellular localization of IL10 via immunofluorescence and immunohistochemistry on D15, D18, D20, D25 and, D40 of pregnancy. IL10 ImR was mostly observed in the endometrial LE and GE, in a pattern dependent on pregnancy stage (Figure 5A). At D15 IL10 ImR was detected in both the LE and GE, but its presence was variable among specimens. Robust IL10 ImR was detected in the LE in D18 samples but was absent in the GE in the same specimens examined (Figure 5B). From D18 to D40 IL10 expression in the LE was consistently observed. In contrast, IL10 ImR in the GE was only detected consistently on D20 (Figure 5C), with variable detection on D25 and D40. The robust detection of IL10 ImR in GE on D20 coincides with a precipitous increase in the abundance of *IL10*-RNA from D15 to D20 (2.7x increase, p=0.0116) and a lower and more variable abundance on D25 (Figure 5F). On D25, some specific IL10 ImR was also observed in the trophoblast layer, and the amniotic membrane (Figure 5D). On D40, strong expression in the LE was detected (Figure 5E). Stromal distribution of IL10 ImR was difficult to assess due to confounding auto-fluorescence and non-specific binding to interstitial blood plasma. It is thus likely that the transition from the pre- to post-attachment phases is associated with a pulse of IL10 secretion from endometrial epithelia.

**Figure 5:**
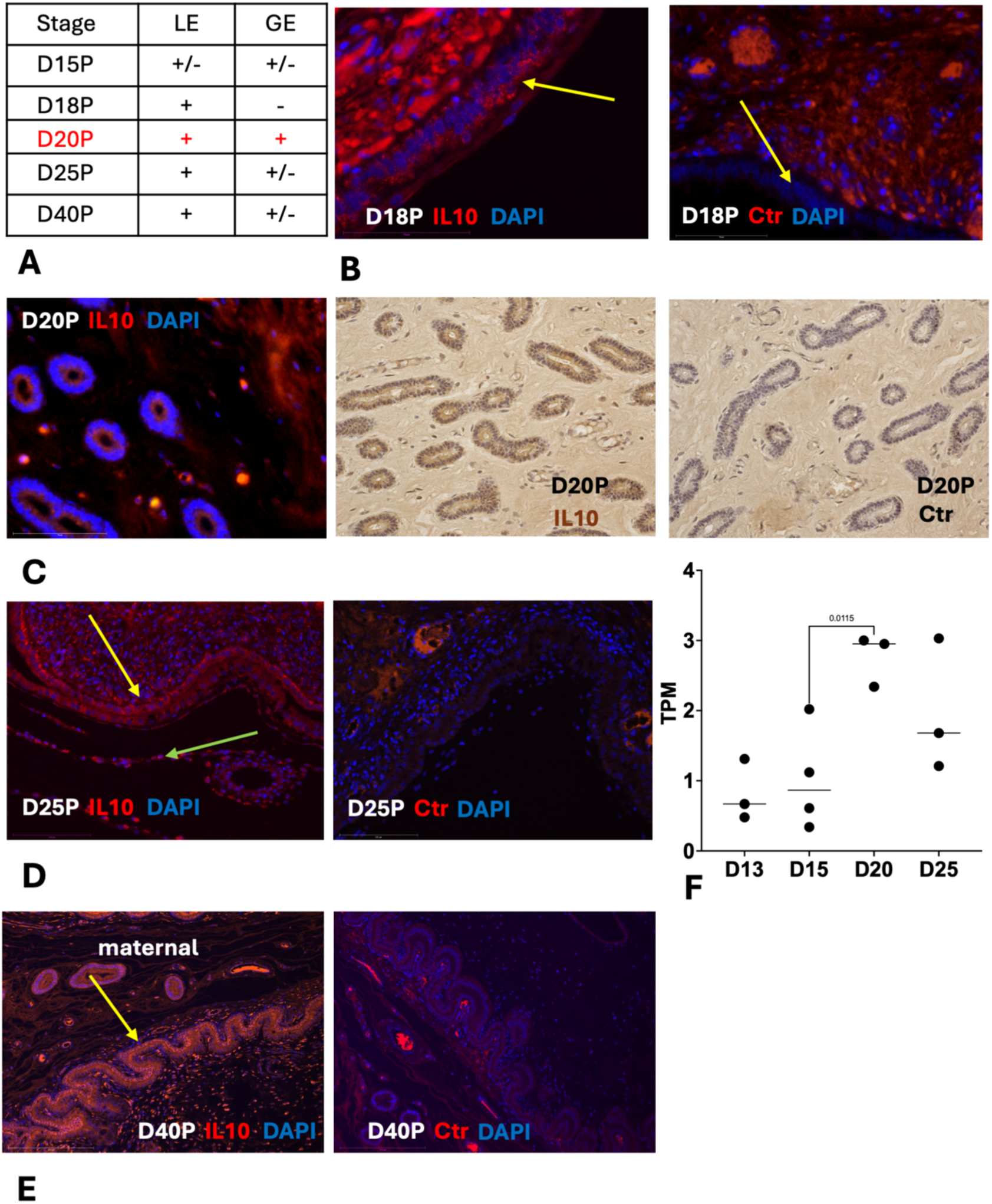
Expression of IL10 in the porcine endometrium: **A**) qualitative summary of observations, + means reliable detection, - not observed, +/- variable detection. **B**) IL10 expression in the LE on D18 of porcine pregnancy. **C**) immunoreactivity for IL10 in the glandular epithelium on D20. **D**) IL10 immunoreactivity in the LE (yellow arrow) and in the allantois (green arrow). **E**) IL10 immunoreactivity in the LE on D40 of the porcine pregnancy. **F**) IL10 RNA expression profile across D13, D15, D20 and D25 of pregnancy. Note the peak in expression on D20 and lower expression on D25.

### Eosinophils are primarily found near the luminal epithelium without attached trophoblast

We investigated the presence and distribution of eosinophils during pre- and post-attachment phases, that is, D13 (n=1), D15 (n=1), D17 (n=1), D18 (n=2), D20 (n=3), D25 (n=3), and D40 (n=3). Eosinophils were observed frequently in the sub-epithelial compartment of the endometrium from D13 to D15 (Suppl. Figure 4A and B), but rarely in D20 to D40 specimens, except in sections in which there was no conceptus. This is consistent with the observations by Kaeoket and collaborators, who reported the highest abundance of eosinophils on D11 of a sampling scheme that includes D1, D3, D11 and, D19 (Kaeoket, Persson et al. 2003). In one section from D13 a mast cell was identified in the sub-epithelial layer of the stroma (Suppl. Figure 4C). This pattern suggests that in the absence of conceptus attachment eosinophils are present in the sub-epithelial layer and close to uterine glands but vacate the stratum compactum compartment after attachment of the trophoblast to the LE suggesting an anti-eosinophil activity of the engaged LE-trophoblast complex. Interestingly, in sheep Monocyte Chemotactic Protein 1 (MCP1) and MCP2 expressing eosinophils increase in the stratum compactum stroma during the attachment phase of implantation in response to progesterone and interferon tau, and are hypothesized to be involved in endometrial remodeling for implantation in that species (Asselin, Johnson et al. 2001).

Comparing these histological observations with expression of mRNAs for known eosinophil chemo-attractants (Rothenberg and Hogan 2006), only Eotaxin 3 (CCL26) and RANTES (CCL5) are expressed at substantial levels. In contrast, Eotaxin 1 and IL5, a cofactor of Eotaxin 1 activity, were detected at very low levels. Both Eotaxin 3 and RANTES have similar expression profiles during the peri-attachment period of pregnancy, with increased average expression from D13 to D20 and lower expression on D25, although the peak average expression of Eotaxin 3 is due to one outlier value (Suppl. Figure 4D and E). This and the lack of Eotaxin1 and IL5 expression suggest that eosinophil recruitment is based on an IL5-independent pathway in the porcine endometrium during the peri-attachment period of pregnancy. This is reminiscent of the fact that IL5 deficient mice still have eosinophils in their endometrium (Robertson, Mau et al. 2000). The highest expression of RANTES on D20, however, coincides with decreasing eosinophil abundance in our results as well as that of (Kaeoket, Persson et al. 2003), suggesting that trophoblast-LE attachment leads to an inhibitory signal or the reduction of a chemoattractive signal(s) for eosinophil recruitment from the LE. In humans, Eotaxin expression is localized to LE and uterine glands (Hornung, Dohrn et al. 2000), and thus regulation of eosinophil recruitment is likely regulated at the level of endometrial epithelia.

## Discussion

The porcine blastocyst establishes itself in the uterus without overt signs of a nidatory injury to the endometrium. Nevertheless, during the peri-attachment period, clear signs of inflammatory and immune activity have been reported, which are resolved after embryo attachment has been established (McLendon, Seo et al. 2020, Johns, Lucas et al. 2021, Sponchiado, Gu et al. 2025). In this study we aimed to gain insight into the mechanisms responsible for the resolution of attachment-associated inflammatory processes during the peri-attachment period of pregnancy in pigs.

As appropriate for this study, the bulk of the following discussion focuses on changes in the inflammatory state of the pig endometrium during the peri-attachment period of pregnancy. But we note that these bulk transcriptomic data complement two previously published reports comparing endometrial transcriptomes from D12 and D14 of pregnancy in pigs with corresponding stages during the non-pregnant cycle (Samborski, Graf et al. 2013, Samborski, Graf et al. 2013). The main findings from these studies, relevant to results of the present study, are increased expression of inflammatory and immune genes in the pre-attachment stages D12 and D14 compared to the non-pregnant uterus, as well as increased abundance of RNAs related to genes responsible for apoptotic processes.

In this study, we documented gene expression throughout the peri-attachment period of pregnancy, from D13 to D25. It is noteworthy that the Spearman correlation data shown in Figure 1A clearly indicate a distinct temporal pattern in endometrial gene expression up to D15, which changes during the transition to D20 of pregnancy. This transition closely reflects previously established changes in the metabolic activity of pig conceptuses, providing confidence that these data fit into what is already understood about this period of pregnancy in the pig. The conceptuses of pigs require energy to support elongation and attachment, and studies have been conducted to elucidate the metabolism of glucose, fructose, and glutamine by elongating pig conceptuses. Although both D15 and D16 pig conceptuses metabolize glucose, fructose, and glutamine, a striking observation was that D15 conceptuses are always significantly more active in their metabolism than those on D16. In addition, several enzymes required for metabolism of glucose, fructose, and glutamine are more highly expressed by conceptuses and endometrium on D13 through D15 than on D16. At publication of these previous studies the authors stated that it appears porcine conceptuses are changing their metabolic strategy between D15 and D17 of gestation and our findings in this study suggest that this transition in the metabolic activity of the conceptuses is mirrored in gene expression of the endometrium (Steinhauser, Landers et al. 2016, Kramer, Steinhauser et al. 2020, Seo, Kramer et al. 2022, Johnson, Seo et al. 2023).

In the present study, all endometrial samples from the same day of pregnancy, i.e., either D13, D15, D20, and D25 are internally consistent in their gene expression profiles. Further, gene expression profiles during D13 and D15 are highly correlated with one another but differs dramatically from gene expression during D20 and D25, which are themselves highly correlated with one another in this study (Figure 1A, B). The exceptions are samples from D17 of gestation. The close association between previously observed changes in the metabolic strategy of elongating and attaching conceptuses and the changes in endometrial gene expression reported here is noteworthy and provides a solid physiological context for the endometrial gene expression data provided in this manuscript.

After wounding and/or infection, resolution of inflammation (RoI) is an active process initiated by the effectors of the initial inflammatory response (Serhan and Savill 2005). It starts with the elimination of the inflammatory stimuli and is followed by a sequence of cellular events leading to the restoration of tissue integrity. These involve 1) the dampening of pro-inflammatory signals and reduction of their receptors, 2) clearing of pro-inflammatory leukocytes such as neutrophils and eosinophils and 3) production of anti-inflammatory signals such as IL10 and TGFb1 stimulating the restoration of tissue integrity. A key event in this sequence is the connection between stage 2, the removal of pro-inflammatory leukocytes, and stage 3, the production of anti-inflammatory and restorative signals. Removal of pro-inflammatory leukocytes can be achieved by reduced recruitment, re-migration of cells into the blood or lymphatic circulations or apoptosis followed by efferocytosis by macrophages. Efferocytosis by macrophages also initiates the alternative activation of macrophages leading to the so-called M2 phenotype, characterized by the secretion of anti-inflammatory signals. To understand the extent to which this model of RoI applies to the quasi-inflammatory stage of pregnancy during conceptus attachment to the endometrial LE, we analyzed our bulk transcriptomic data with samples from D13 to D25 of pregnancy and supplemented these results with immuno-localization to identify the cellular sources of some of the key transcripts.

Our data show that the rate of endometrial transcriptome change is high up to D20 and then slows. This transition coincides with the establishment of firm trophoblast attachment to the LE and reduction in the presence of eosinophils within the endometrium. The transition from D15 to D20 also coincides with an increase in the level of expression of RNAs for genes expected to be involved in RoI. These include significant upregulation of the prostaglandin catabolic enzyme, *HPGD* (Figure 2C), an enzyme that may be involved in decreasing prostaglandin signaling after implantation (Hansen, Keelan et al. 1999), consistent with the results of Seo and colleagues who reported a precipitous increase of *HPGD* RNA from D15 to D30 and increasing expression up to D90 (Seo, Choi et al. 2014). In addition, we found increased expression of genes generically associated with M2 macrophage polarization: CD86, a cell surface receptor of antigen-presenting cells such as dendritic cells and macrophages; MRC1 (CD206), a mannose receptor involved in clearing pro-inflammatory chemokines and assisting in efferocytosis; and ARG1, a marker of M2 macrophages. ARG1 is responsible for the conversion of arginine into ornithine, where the latter is processed to form either proline or polyamines. Polyamines play key roles in developing and maintaining anti-inflammatory environments (Bjelakovic, Stojanovic et al. 2010). Most notable is a peak in *IL10* RNA abundance at D20, followed by continued enhanced expression on D25. IL10 is an important anti-inflammatory signal that negatively regulates a broad array of pro-inflammatory cytokines (Ouyang, Rutz et al. 2011). It is generically expressed in M2 macrophages as well as type 2 T-helper cells, Th2. Overall, there is a consistent pattern of transition from a pro-inflammatory environment prior to conceptus attachment documented previously (Samborski, Graf et al. 2013, McLendon, Seo et al. 2020), to a regime typical for the resolution of inflammation post-attachment. Immuno-localization of some of these anti-inflammatory mediators, however, suggests different cellular origins for a majority of these molecules than would be expected in a generic RoI.

ARG1 transcripts are virtually absent in pre-attachment samples and are only moderately present starting at D20 (Figure 3A). Immuno-reactivity was found in large round cells resembling macrophages in the sub-epithelial stratum compactum stroma of the endometrium (Figure 3F).

MRC1/CD206 ImR was found in two compartments. One was the endometrial stroma, identifying cells with fibroblast-like morphology and a few macrophage-like cells (Figures 3B, C, and E). In addition, macrophage-like MRC1-positive cells were located in the placental compartment (Figure 3D). Cells resembling the MRC1-positive cells in the placental compartment are positive for osteopontin (Wing, Erikson et al. 2020), but a formal definition of these cells has, to our knowledge, not been achieved, but could be Hofbauer cells.

The increase in MRC1 positive cells after attachment is also interesting as MRC1 is known to facilitate clearance of cytokines and interleukins that are decorated with mannose containing glycans. Human IFNG has been shown to be glycosylated on Ans-97 with a mannose rich glycan (Sareneva, Mortz et al. 1996) and thus porcine IFNG could be cleared from the endometrium by MRC1 expressing stromal and immune cells.

*IL10* has moderate RNA abundance in the endometrium prior to D20, followed by a peak of expression on D20 and then lower but still elevated expression on D25 (Figure 5F). IL10 ImR was found primarily in the LE from D13 to D40, and IL10 protein was transiently but strongly expressed on D20 in the endometrial GE (Figure 5C), coincidental with the peak of *IL10* RNA expression in the bulk transcriptome. Our findings are consistent with those of Han and collaborators, who reported expression of IL10 in the LE on D15 and D40, but that group did not examine endometrial samples from D20 or D25 (Han, Yoo et al. 2022). Han et al. also reported ImR in rare cells in the stoma, which they interpreted to be putative macrophages, a finding we could not replicate.

CD86 is a cell surface ligand on antigen-presenting cells, including macrophages. Together with CD80 this molecule engages with receptors on T-cells to either activate naïve T-cells or to inhibit T-cell activity, depending on the kind of receptor the T-cell is displaying (CD28 for activation and CTLA4/CD152 for inhibition). In our preparations, we found CD86 ImR at the apical surface of trophoblast cells attached to LE on D20. On D25 CD86 ImR localized to the apical cytoplasm of LE, a pattern reminiscent of trogocytosis that is reported to occur during interactions between macrophages and Tcells (Joly and Hudrisier 2003). Trogocytosis is the incorporation of a ligand-receptor complex in one of the two interacting cell types, usually T-cells interacting with macrophages. *CTLA4* and *CD28* transcripts, the two generic ligands for CD86, were present in our RNAseq data, but at very low levels of abundance. Attempts to localize CTLA/CD152 and CD28 ImR in the LE were unsuccessful, but on D20 we observed some putative intra-epithelial lymphocytes in the GE expressing CD152 (Figure 4B). The potential role of CD86 in the trophoblast and its attachment to the LE is rather enigmatic. It is likely a part of trophoblast to LE signaling, but whether our observations indicate a role of the trophoblast in presenting conceptus antigens to the LE, is an intriguing possibility, but as yet untested.

Overall, the pattern arising from our study is that the transition to firm trophoblast-LE attachment is associated with expression of mediators often involved in the resolution of inflammation. The disappearance of eosinophils from this environment is consistent with this observation (Suppl. Figure 3). The surprising finding, however, is that the cellular sources of some of these molecules are different than predicted from the generic RoI model (Serhan and Savill 2005, Sugimoto, Sousa et al. 2016, Panigrahy, Gilligan et al. 2021). CD86 is found in the trophoblast, MRC1 on putative fibroblasts and IL10 in the LE and GE, with ARG1 the only M2 macrophage marker found in putative macrophages in the subepithelial layer of the endometrial stroma. This pattern suggests that the overall mechanism of RoI seems to be the same as that after infection or wounding in the rest of the body, but at the porcine uterine-placental interface it is mediated by different cell types than in other tissues. This observation is coherent with the fact that in the pig the vast majority of IFNG within the intrauterine environment is produced by the trophoblast instead of M1 macrophages or other immune cells (Joyce, Burghardt et al. 2007). In contrast, during mouse implantation IFNG is produced by uterine natural killer cells (Croy, He et al. 2003).

An intriguing aspect of porcine fetal-maternal communication is the role of IFNG produced by the trophoblast. IFNG is generally considered to be a pro-inflammatory signal, resulting in activation of inflammatory macrophages. Interestingly, gilts receiving IFNG -/- blastocysts develop what appears to be a runaway uterine inflammation and loss of pregnancy (Johns, Lucas et al. 2021). A single-nuclei study showed that at D15, the lack of IFNG signaling from the blastocyst affects the cell type composition in the endometrium (Sponchiado, Gu et al. 2025). Specifically, the authors observed a strongly reduced abundance of macrophages and osteopontin-positive LE cells. One likely interpretation is that trophoblast-derived IFNG is necessary for the recruitment of macrophages and, failing that, the potential for an M2 mediated resolution of inflammation is lacking leading to the unchecked inflammation associated with IFNG -/- blastocysts. Our results, however, do not support an important role of M2 activation in resolution of pregnancy, as the main cellular sources of IL10 are epithelia consistent with the results of Han and collaborators (Han, Yoo et al. 2022). However, the results of the single-nuclei study suggest an alternative pathway for the effects of IFNG -/- embryos, namely the lack of osteopontin (OPN, a.k.a. Secreted Phosphoprotein 1, SPP1) in LE. This observation in conjunction with the findings in this study suggest a model for why blastocyst IFNG signaling is necessary for resolution of inflammation in the porcine endometrium (Figure 6).

**Figure 6:**
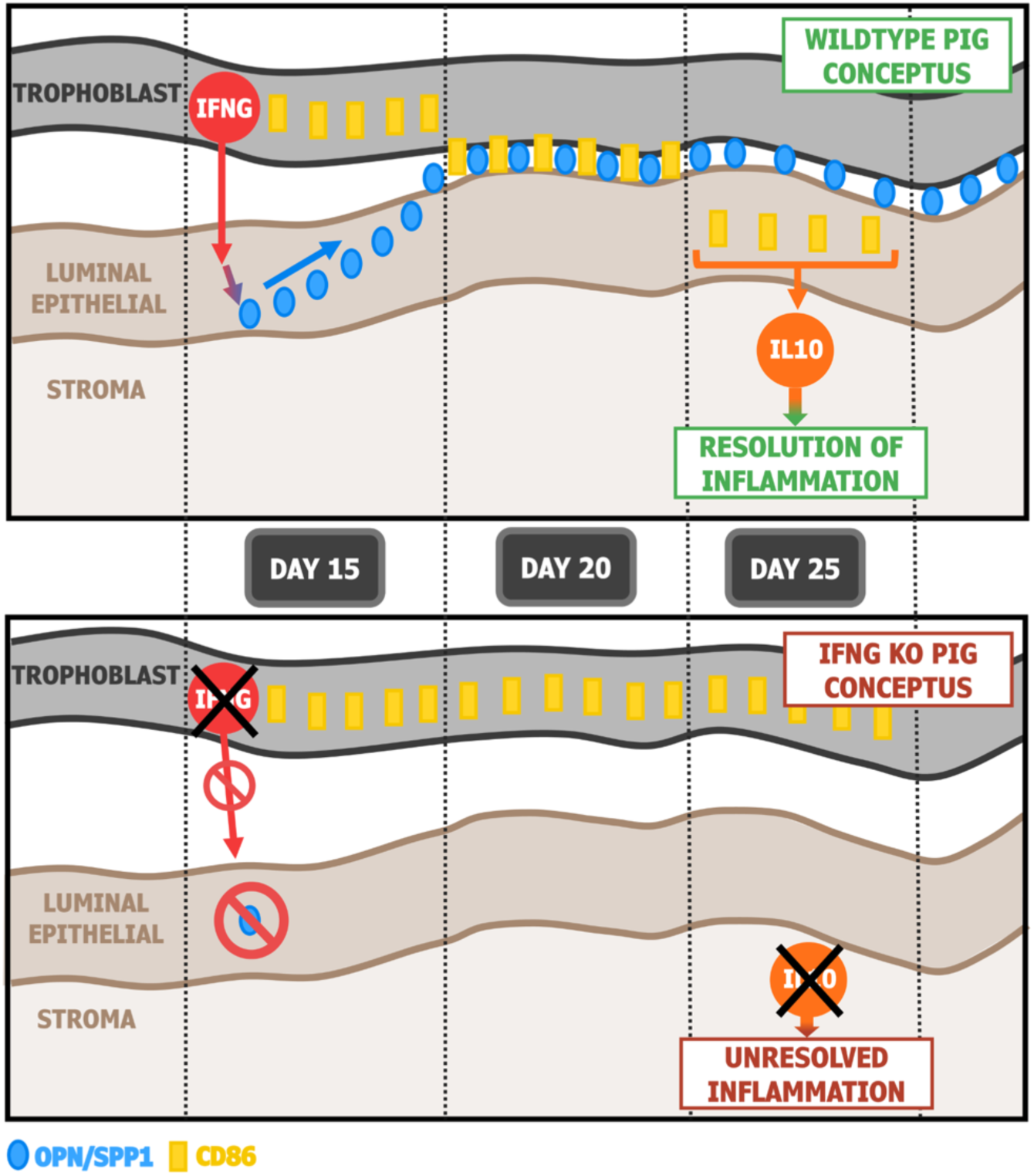
Model of how IFNG signaling from the trophoblast may contribute to the resolution of inflammation in the porcine peri-attachment endometrium. **Top image**: on D15 IFNG from the trophoblast stimulates the production of OPN/SSP1 in the luminal epithelium. At the same time CD86 protein is accumulating in the trophoblast cells. On D20 the OPN is exported from the LE providing the molecular bridge for the blastocyst attachment to the LE. Once the contact is established, CD86 is inserted in the apical cell membrane of the trophoblast and engages with an unknown receptor on the LE side. By D25, the CD86 – receptor complex is transferred to the LE cells where CD86 immunoreactivity is found in their apical cytoplasm vial a process akin to trogocytosis. This model assumes that this transfer is instrumental in inducing an increase in IL10 expression in epithelial cells and thus contributes to the resolution of inflammatory processes in the endometrial stroma. **Bottom image**: in the absence of blastocyst IFNG the LE does not produce OPN/SPP1 and thus the trophoblast cannot attach to the LE, also preventing the engagement of CD86 with a hypothetical receptor on the LE. According to the model, this also prevents the production of IL10 in the endometrial epithelia and the stromal inflammation proceeds unopposed as observed in the IFNG KO experiment of Johns et al. (2021). In fact, these authors also report a failure of attachment of IFNG -/- blastocysts and Sponchiado and colleagues (2025) reported a lack of SPP1 producing LE cells in the presence of IFNG -/-blastocysts on D15.

SPP1 is abundantly expressed by the endometrial LE during the peri-attachment phase of pregnancy in pigs (Garlow, Ka et al. 2002, White, Ross et al. 2005). SPP1 and its integrin receptors are proposed to be molecular building blocks of the trophoblast-LE attachment complex (Erikson, Burghardt et al. 2009, Frank, Seo et al. 2017). If the lack of IFNG leads to a lack of SPP1 production by LE, as shown by Sponchiado and collaborators (Sponchiado, Gu et al. 2025), it can be inferred that it negatively affects attachment of the trophoblast to the LE. Failure of proper attachment may further prevent necessary signal transduction that depends on direct contact between the trophoblast and the LE. A case in point would be the formation of integrin adhesion complexes (IACs) that assemble in both trophoblast and LE cells during conceptus attachment in pigs (Johnson, Burghardt et al. 2023, Johnson, Minela et al. 2025). Another potential interaction could be CD86-mediated signaling between trophoblast and LE (Figure 4A). If the CD86 mediated trophoblast-LE interaction has a role in RoI, which is not yet proven, the loss of expression of SPP1 by LE could prevent epithelial IL10 expression and thus compromise the establishment of pregnancy (Figure 6).

From an evolutionary point of view, it is interesting to note that the epitheliochorial placentation of pigs and other artiodactyls is derived from a hemochorial condition (Elliot and Crespi 2009). In hemochorial species, the control of nidatory inflammation is performed by the interaction of decidual cells with uterine leukocytes, such as Treg cells and uterine macrophages, as the most likely mediators (Erlebacher 2013, Chavan, Griffith et al. 2021). However, we are not aware of a targeted study of the resolution of nidatory inflammation in hemochorial species. In any case, the pig and other artiodactyls do not have decidual cells, which usually develop in response to implantation of blastocysts, except for a minority of species, such as apes including humans, which evolved spontaneous decidualization (Critchley, Babayev et al. 2020). From these comparative facts one may infer that the evolution of epitheliochorial placentation entailed the loss of decidual cells due to the loss of invasive implantation by blastocysts. As a consequence, control of the uterine immune environment, normally regulated by decidual cells, was likely transferred to other cell types present within the uterine-placental environment including epithelia and fibroblasts that maintained the production of immune regulatory molecules such as IL10 and others.

At a general level, the expression of immune regulatory molecules by non-immune cells in the uterus during pregnancy can be understood as a means of decoupling immune regulation in the pregnant uterus from that in the rest of the body. Such a decoupling may achieve pregnancy-related immune suppression at the fetal-maternal interface but retains active immune defenses in the rest of the body, protecting the health of the mother during pregnancy.

While our results suggest an active process of resolution of endometrial inflammation involving non-immune cells in the porcine endometrium, they also raise a number of new questions that the present study is unable to answer. Among them are

- What is the role of CD86 expression by the trophoblast and its putative interaction with the LE, including an apparent trogocytosis?
- Is epithelial IL10 secretion paracrine, i.e., directed towards the stroma, or exocrine, i.e., directed towards the blastocyst? A role of IL10 in conceptus development is suggested in its immunoreactivity in extraembryonic membranes on D25 of pregnancy (Figure 5D).
- What are the signals that control the epithelial expression of IL10 in the endometrium at the time of conceptus attachment?
- How does the IFNG signal from trophectoderm regulate the endometrial expression of anti-inflammatory mediators, and thus how does IFNG participate in the resolution of attachment-associated endometrial inflammation?
- How does LE engagement with the trophoblast regulate eosinophil abundance in the sub-epithelial stroma of the endometrium?
- What is the nature of the MRC1-positive cells in the placenta?

The main limitation of the current study is the lack of single-cell transcriptomic data covering the peri-attachment period. There are single-nuclei data available for D15 (Sponchiado, Gu et al. 2025) and single-cell transcriptome data for D9 and D12 (Zang, Gu et al. 2024). In our study we used bulk RNAseq to probe gene expression dynamics during the entire peri-attachment period and to identify periods of pregnancy where a more detailed investigation is warranted. Based on our results, an in-depth comparison of D15, D20, and D25 of pregnancy would be most informative for understanding the resolution of attachment-associated inflammation.

## Materials and Methods

### Animals and Tissue Collection

Sexually mature, 8-month-old female crossbred pigs (gilts) were observed daily for estrus (D0) and exhibited at least two estrous cycles of normal duration (18 to 21 days) before being used in these studies. All experimental and surgical procedures were in compliance with the Guide for Care and Use of Agricultural Animals in Teaching and Research and approved by the Institutional Animal Care and Use Committee of Texas A&M University.

Gilts were bred naturally to boars with proven fertility and were euthanized and then ovariohysterectomized on either day 13 (n = 3), 15 (n = 3), 17 (n = 2), 20 (n = 3), 25 (n = 3), or 40 (n = 4) of pregnancy. Tissue sections (uterus or uterus with attached placenta ∼1 cm thick) from the middle of each uterine horn of all hysterectomized gilts were fixed in fresh 4% paraformaldehyde in PBS (pH 7.2) and embedded in Paraplast-Plus (Oxford Laboratory, St. Louis, MO) for immunohistochemistry. The remaining endometrium was physically dissected from the myometrium, snap-frozen in liquid nitrogen, and stored at -80°C for RNA extraction.

### RNA Extraction and Sequencing

Total RNA was extracted from endometrial and placental tissue samples using Trizol reagent (Life Technologies, Carlsbad, CA, Cat#15596018) and RNeasy Mini Kit (Qiagen, Cat#74104) according to the manufacturer’s recommendations. Quality and concentrations of RNA samples were determined using a NanoDrop Spectrophotometer (ND-1000; ThermoFisher Scientific). Samples were sent to the Texas A&M Molecular Genomics Core for Tapestation QC (TapeStation Analysis Software 5.1) and RNA sequencing (P2-XLEAP - 200 cycle).

Raw data were analyzed by the University of Vienna, Austria. The Trim-Galore version 0.6.10 with default parameters in paired-end mode was used to trim the reads. STAR version 2.7.11a was used for mapping the reads on the Sscrofa 11.1 assembly with the gene annotation from Ensembl release 110. Building the index and mapping the reads was done with default parameters. Subsequently, featureCounts v2.0.6 was used with the -p and --countReadPairs parameters to count the number of read pairs per gene. The list of gene orthologs between pig and human was downloaded from BioMart to supplement missing gene names in the pig gene annotation. The output files from featureCounts were reformatted, gene names extended and TPM calculated using custom python scripts.

To quantify the total amount of transcriptome change we calculate the angle between the gene expression vectors from consecutive samples. Each sample type is represented as a vector of average gene expression levels measured in TPM, say for one type of sample the gene expression profiles is

**x**=(x_1_, …, x_n_)

where the n is the number of genes included in the expression profile, and

**y**=(y_1_, …, y_n_) for the other sample type.

The similarity between these two vectors can be represented by the angle of these vectors in the expression space and can be calculated from the inner product of the two vectors

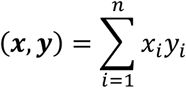

From which the angle can be calculated as

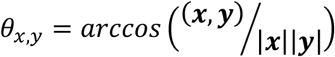

Where |***x***| and |***y***| are the Euclidian length of the two expression vectors.

The angle *θ_x,y_* is a measure of divergence between **x** and **y**. In order to estimate the rate of change we need to normalize the difference with the amount of time that separates the two samples. We used two ways to normalize for the amount of time between the sample: 1) divide by the number of days between the sampling times and 2) divide by the square root of the number of days. Method 1 assumes that changes from day to day happen in a straight line within in the expression space. Method 2 assumes that the direction of change from day to day is random, which in high-dimensional space converges toward orthogonal change. This predicts that the Euclidian distances are on average the distance divided by the square root of the number of days.

If two vectors with TPM values are of different length, it means that the expression profile of the longer vector has a steeper profile, i.e., the expression levels are more uneven than in the shorter vector. The reason is that the sum of the TPM vectors is always 10^6^, per definition, and thus the Euclidian length of the vector is proportional to the variance of the coefficients in the vector. Hence a longer vector of TPM values has a steeper expression profile than a TPM vector with smaller length.

### Immunofluorescence Analysis

Immunofluorescence analyses for immune cell markers was performed on paraffin-embedded tissues from gilts on D13, 15, 17, 20, and 25 of pregnancy. Paraffin-embedded tissues were serially sectioned at 5 µm thickness using a Leica RM2235 Microtome and fixed to a microscope slide. The sections were then deparaffinized with Citri-Solv (Fisherbrand, Cat#22143975) and rehydrated in decreasing concentrations of ethanol (100%, 95%, 70%, DI H2O). Boiling citrate was used for antigen retrieval. Sections were then washed in 1X PBS and blocked with 10% normal goat serum at room temperature for 1hr before applying the primary antibodies (rabbit anti-myeloperoxidase antibody, Abcam, Cat#ab9535, 1:50; mouse anti-TLR4 antibody, Abcam, Cat#ab22048, 1:50; mouse anti-liver arginase antibody, Abcam, Cat#ab239731, 1:200) or the negative control (rabbit IgG, EMD Millipore, Cat#12-370; mouse IgG, EMD Millipore, Cat#12-371) at equal concentrations. Slides were incubated overnight at 4℃. After overnight incubation, sections were washed in 1X PBS and incubated in a 1:250 dilution of the secondary antibody (goat anti-rabbit Alexa Fluor 488, Life Technologies, Cat#A-11034; goat anti-rabbit Alexa Fluor 594, Life Technologies, Cat#A-11012; goat anti-mouse Alexa Fluor 488, Life Technologies, Cat#A-21042) at room temperature for 1hr. The slides were then counterstained with Prolong Gold Antifade reagent with DAPI (Cat#P36935) and coverslips were applied. Images were captured with an Axio Imager.M2 microscope (Carl Zeiss, Thornwood, NY, USA) interfaced with an Axioplan HR digital camera.

### Immunohistochemical Analysis

Immunohistochemical analyses for macrophage markers was performed on paraffin-embedded tissues from gilts on D15, D20, and D25 of pregnancy. Paraffin-embedded tissues were serially sectioned at 5 µm thickness using a Leica RM2235 Microtome and fixed to a microscope slide. The sections were then deparaffinized with Citri-Solv (Fisherbrand, Cat#22143975) and rehydrated in decreasing concentrations of ethanol (100%, 95%, 70%, DI H2O). Boiling citrate was used for the antigen retrieval. Sections were then set in 200mL methanol and 6.7mL of H_2_O_2_ for 15 minutes, followed by 2 rinses in DI water. Sections were then washed in 1X PBS and blocked with TBS-T (Triton-X, Tris) at room temperature for 1hr. Primary antibodies (rabbit anti-CD206 antibody, Proteintech, Cat#81525-1-RR, 1:300; rabbit anti-CD86 antibody, Abbiotec, Cat#251407, 1:300) or the negative control (rabbit IgG, EMD Millipore, Cat#12-370) at equal concentrations were applied and incubated overnight at 4℃. Sections were then washed in 1X PBS and secondary antibody (goat anti-rabbit IgG H&L HRP, Abcam, Cat#ab6721, 1:1000) for 1hr at room temperature. After washing with 1X PBS, sections were incubated in DAB for 10 minutes and dunked in DI water. Sections were stained with hematoxylin and rinsed in DI water, followed by dehydration in ascending concentrations of ethanol and Citrosolv. The slides were then counterstained with Toluene reagent (Cat#) and coverslips were applied. Images were captured with an Axio Imager.M2 microscope (Carl Zeiss, Thornwood, NY, USA) interfaced with an Axioplan HR digital camera.

Immunohistochemical analyses for IL-10 was performed on paraffin embedded tissues from gilts on D15, D20, and D25 of pregnancy. Paraffin embedded tissues were serially sectioned at 5 µm thickness using a Leica RM2235 Microtome and fixed to a microscope slide. The sections were then deparaffinized with Citri-Solv (Fisherbrand, Cat#22143975) and rehydrated in decreasing concentrations of ethanol (100%, 95%, 70%, DI water). Following the dips in DI water, sections were then washed in 1X PBS and blocked with TBS-T (Triton-X, Tris) at room temperature for 1hr as no antigen retrieval method was used. Sections were then set in 200mL Methanol and 6.7mL of H2O2 for 15 minutes, followed by 2 rinses in DI water. Primary antibodies (goat anti-IL10 antibody, Bio-Techne, Cat# AF693, 1:300) or the negative control (TBS-T) were applied and incubated overnight at 4℃. Sections were then washed in 1X PBS and secondary antibody (donkey anti-goat IgG H&L HRP, Abcam, Cat# ab97110, 1:500) for 1hr at room temperature. After washing with 1X PBS, sections were incubated in DAB for 10 minutes and dunked in DI water. Sections were stained with hematoxylin and rinsed in DI water, followed by dehydration in ascending concentrations of ethanol and Citrosolv. The slides were then counterstained with Toluene reagent (Cat#) and coverslips were applied. Images were captured with an Axio Imager.M2 microscope (Carl Zeiss, Thornwood, NY, USA) interfaced with an Axioplan HR digital camera.

### Eosinophil and Mast Cell detection

In order to locate eosinophil and mast cells we stained deparaffinized histological sections with the Abcam hematological staining kit (ab150665) according to manufacturer’s instructions and examined the slides under transmission light microscope.

## Author Contributions

Conceptualization and study design: FWB, GAJ, GPW,

Lab Work: GPW, TM, AR

Data Analysis: GPW, JE

Funding: FWB, GAJ

## Acknowledgements

the support of the Hagler Institute of Advanced Study at Texas A&M University through a Hagler Fellowship to GPW and a graduate fellowship to Alex Ross is gratefully acknowledged. This support made this collaboration possible. The authors thank the members of the Johnson laboratory for efforts regarding animal husbandry and collection of porcine endometrial tissues. The transcriptomic data was generated at the TAMU Genomics and Bioinformatics service and analyzed at the University of Vienna Life Science Computing cluster. The authors thank Professor Mihaela Pavlicev and her lab members for input and discussion on a previous version of this manuscript.

## Supplemental Figures

**Suppl. Figure 1:**
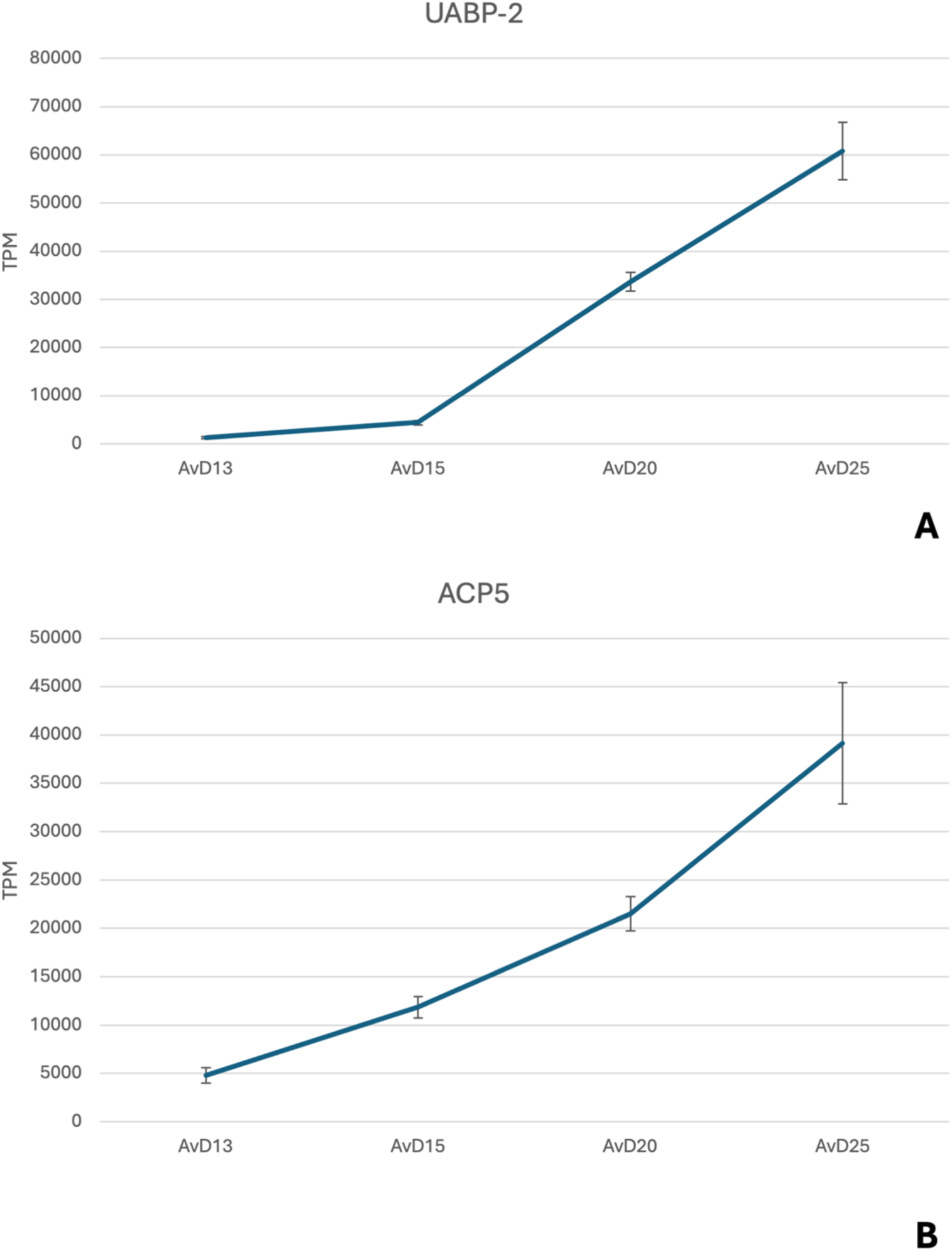
Relative expression of mRNAs is dynamic for *UABP-2* and *ACP5* in the porcine endometrium from D13 to D25.

**Suppl. Figure 2:**
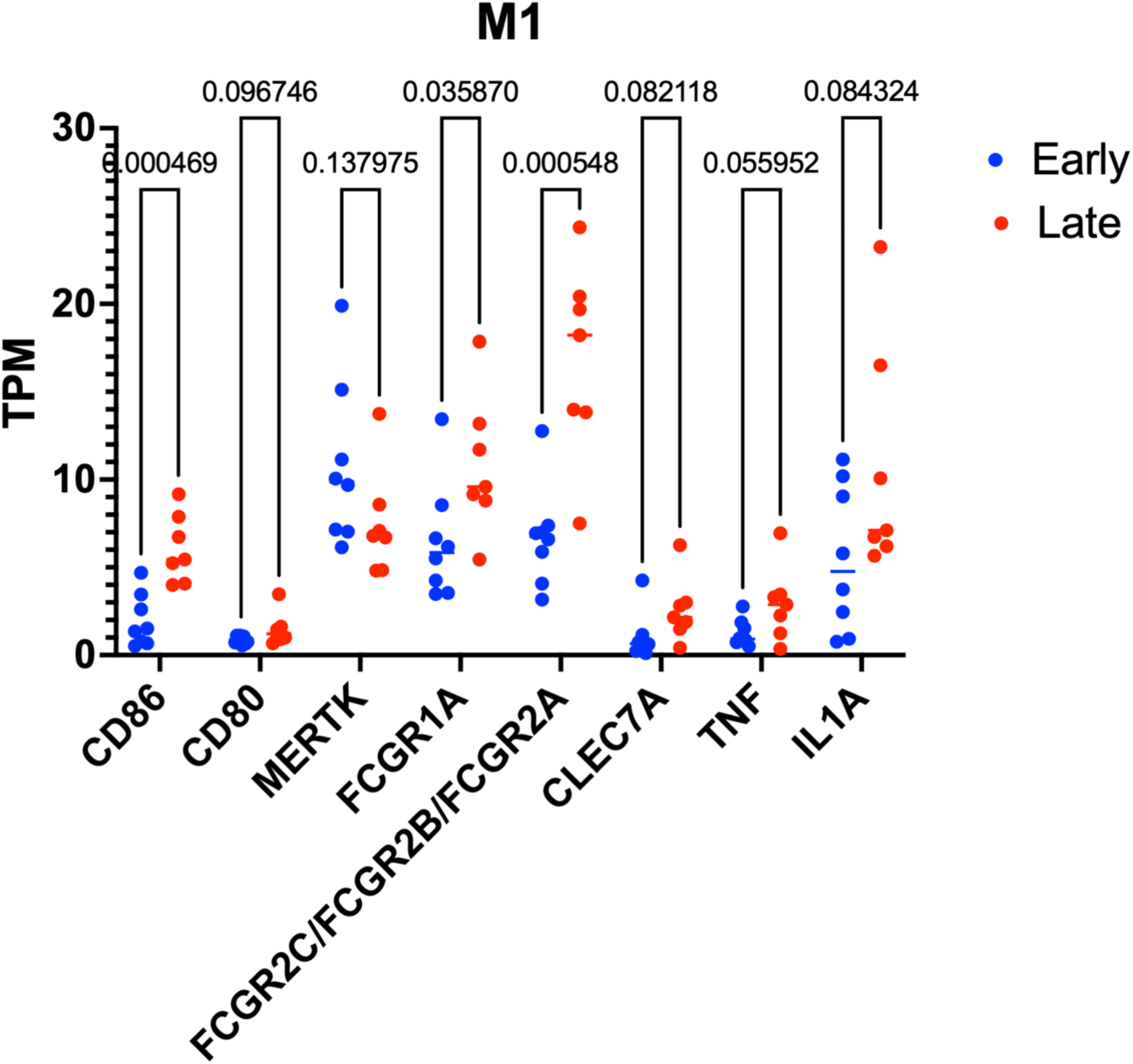
A comparison of the expression of M1 marker genes in the porcine endometrium between pre-attachment and post-attachment periods of pregnancy.

**Suppl. Figure 3:**
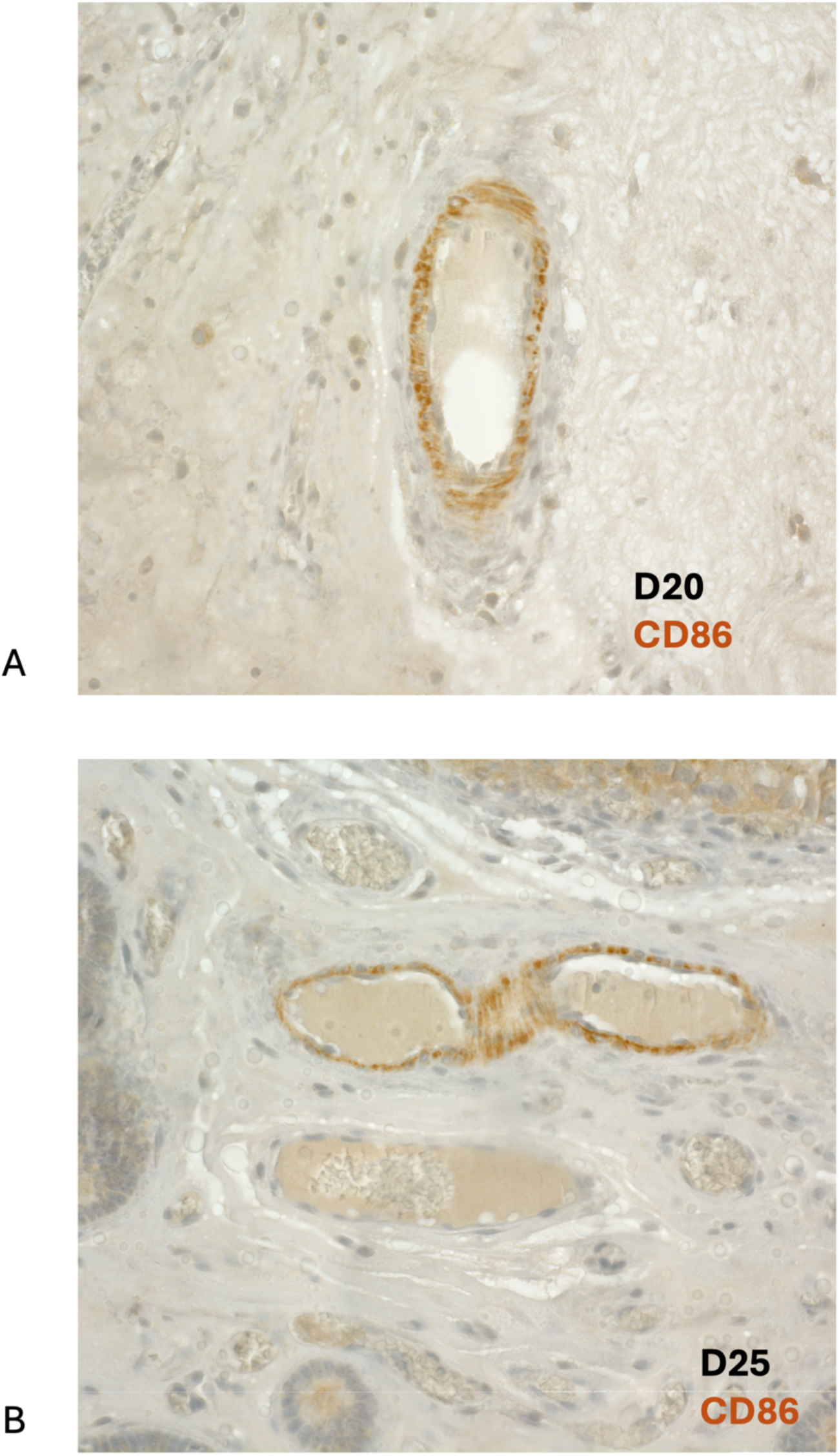
CD86 immunoreactivity in the perivascular cells of the porcine endometrium on D20 (A) and D25 (B).

**Suppl. Figure 4:**
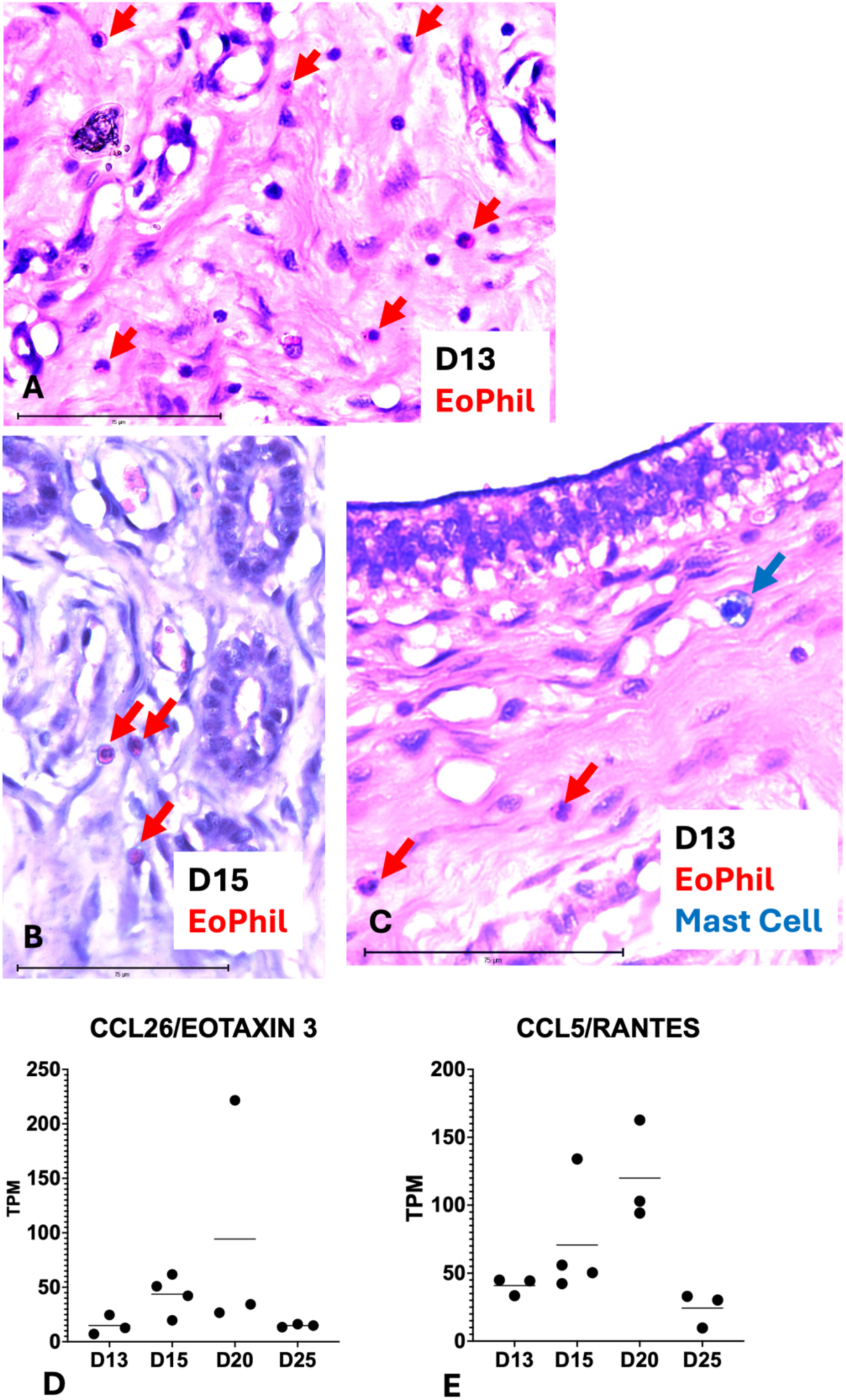
Eosinophils (red arrows) and mast cell (blue arrow) **A**) Eosinophil cells in the porcine endometrium on D13 of pregnancy. **B**) Eosinophil cells in the porcine endometrium on D15 of pregnancy near uterine glands. **C**) A mast cell in the sub-epithelial stroma of the porcine endometrium on D13 of pregnancy. **D** and **E**) RNA expression profile of *Eotaxin3* and *RANTES,* respectively, during the peri-attachment period.

